# Mapping eye, arm, and reward information in frontal motor cortices using electrocorticography in non-human primates

**DOI:** 10.1101/2024.08.13.607846

**Authors:** Tomohiro Ouchi, Leo R. Scholl, Pavithra Rajeswaran, Ryan A. Canfield, Lydia I. Smith, Amy L. Orsborn

## Abstract

Goal-directed reaches give rise to dynamic neural activity across the brain as we move our eyes and arms, and process outcomes. High spatiotemporal resolution mapping of multiple cortical areas will improve our understanding of how these neural computations are spatially and temporally distributed across the brain. In this study, we used micro-electrocorticography (µECoG) recordings in two male monkeys performing visually guided reaches to map information related to eye movements, arm movements, and receiving rewards over a 1.37 cm^2^ area of frontal motor cortices (primary motor cortex, premotor cortex, frontal eye field, and dorsolateral pre-frontal cortex). Time-frequency and decoding analyses revealed that eye and arm movement information shifts across brain regions during a reach, likely reflecting shifts from planning to execution. We then used phase-based analyses to reveal potential overlaps of eye and arm information. We found that arm movement decoding performance was impacted by task-irrelevant eye movements, consistent with the presence of intermixed eye and arm information across much of motor cortices. Phase-based analyses also identified reward-related activity primarily around the principal sulcus in the pre-frontal cortex as well as near the arcuate sulcus in the premotor cortex. Our results demonstrate µECoG’s strengths for functional mapping and provide further detail on the spatial distribution of eye, arm, and reward information processing distributed across frontal cortices during reaching. These insights advance our understanding of the overlapping neural computations underlying coordinated movements and reveal opportunities to leverage these signals to enhance future brain-computer interfaces.

Significance statement

Picking up your coffee mug requires coordinating movements of your eyes and hand and processing the outcomes of those movements. Mapping how neural activity relates to different functions helps us understand how the brain performs these computations. Many mapping techniques have limited spatial or temporal resolution, restricting our ability to dissect computations that overlap closely in space and time. We used micro-electrocorticography recordings to map neural activity across multiple cortical areas while monkeys made goal-directed reaches. These measurements revealed high spatial and temporal resolution maps of neural activity related to eye, arm, and reward information processing. These maps reveal overlapping neural computations underlying movement and open opportunities to use eye and reward information to improve therapies to restore motor function.

## Introduction

Reaching involves coordinating our limbs and eyes (Crawford et al., 2004; Song and McPeek, 2009; Vazquez et al., 2017). When picking up a mug, one typically looks at it first. Goal-directed movements also incorporate feedback like rewards. If the mug spills, one might move differently next time. Each reach gives rise to neural activity distributed across brain areas as we process eye and arm movements and outcomes.

Several pre-central cortical areas have been implicated in goal-directed reaching. In motor cortex (MC), primary motor (M1) and pre-motor (PM) cortices contribute to reach preparation and execution and their neural activity contains information about movement direction (Tanji and Evarts, 1976; Georgopoulos et al., 1982; Crammond and Kalaska, 2000). In pre-frontal cortices (PFC), the frontal eye field (FEF) and dorsolateral pre-frontal cortex (DLPFC) contain activity related to saccade initiation and direction (Bruce and Goldberg, 1985; Bruce et al., 1985; Funahashi et al., 1990, 1991; Schall, 2002; Everling and DeSouza, 2005; Markowitz et al., 2011; Funahashi, 2014; Boulay et al., 2016; Lee et al., 2017). FEF and DLPFC also exhibit reward-related activity (Roesch and Olson 2003).

Parsing computations within pre-central cortical areas is challenging because information is distributed. Many areas lack discrete functional boundaries. PM appears to contain a gradient of task information, with visual or eye-related activity located rostrally near the AS, while more caudal regions are primarily active during reach execution (Johnson et al., 1996; Fujii et al., 1998, 2000; Cisek and Kalaska, 2005; Nakayama et al., 2016). An area may also perform multiple computations. PM and M1 have reward-(Ramkumar et al., 2016; Ramakrishnan et al., 2017) and gaze-related activity (Pesaran et al., 2006, 2010; Batista et al., 2007). PFC shows activity related to both saccades and reward (Roesch and Olson 2003). Despite this heterogeneity, frontal cortices have often been studied in isolation with sparse sampling of neural activity within regions.

Dissecting the functional properties of frontal cortices is also challenging because eye, arm, and reward computations do not operate in isolation. For example, coordinating a reach with a saccade improves reach accuracy independent of vision (Vazquez et al., 2017). Some neurons in PM only fire when eye and arm movements co-occur (Kurata, 2017). Rewards also influence movement parameters (speed, vigor) (Takikawa et al., 2002; Manohar et al., 2015; Summerside et al., 2018). However, studies often use tasks and analyses to study eye, arm, and reward computations in partial isolation.

Functional maps may help improve our understanding of the coordinated computations involved in goal-directed reaching. Functional mapping requires simultaneous recordings across multiple areas paired with behavioral tasks and analyses that explore arm, eye, and reward processing. Imaging modalities like fMRI offer high spatial resolution, but may lack sufficient temporal resolution to distinguish closely timed eye and arm movements (Glover, 2011; Sejnowski et al., 2014; Chehade and Gharbawie, 2023). Neuron-resolution electrophysiology offers high spatial and temporal precision, but current limits on the number of simultaneous measurements constrain spatial coverage. Meso-scale measurements like electrocorticography (ECoG) provide millimeter spatial resolution and can resolve temporal dynamics comparable to neuronal spiking across large areas (Chiang et al., 2020; Trumpis et al., 2021). ECoG has proven valuable to map activity related to reaching or saccades (Ball et al., 2009; Lee et al., 2017; Kaiju et al., 2021), but has not been used to dissect eye, arm, and reward information within frontal cortices.

We used high-density micro-electrocorticography (µECoG) to map eye, arm, and reward related neural signals within PFC and MC in monkeys performing a visually guided reaching task with minimally or unconstrained gaze. We find that µECoG signals captured neural activity related to eye and arm movements and receiving rewards. µECoG maps captured shifts in information about movement across brain regions over time, identified overlapping information about eye and arm movements, and identified reward-related activity. Our results improve our understanding of the heterogeneous functions in PFC and MC while also identifying opportunities and challenges for brain-computer interface applications.

## Materials and Methods

### Surgical procedures

All procedures were approved by the Institutional Animal Care and Use Committee at the University of Washington. Two male rhesus macaques (Monkey 1: Beignet, 9 years old, 11.3 kg; monkey 2: Affogato, 10 years old, 10.6 kg) were implanted with a custom titanium chamber in the left hemisphere as well as a headpost for head fixation (Rogue Research, Montréal, CA). The chambers were stereotaxically targeted based on each animal’s MRI to span portions of primary motor cortex (M1), premotor cortex (PM), frontal eye fields (FEF), and dorsolateral prefrontal cortex (DLPFC). Chambers were also custom shaped to the skull curvature based on MRI data. The implants were attached using self-tapping titanium screws into bore holes made using a handheld drill bit (Veterinary Orthopedic Implants, St. Augustine FL). A 2.6 cm diameter craniotomy was subsequently carried out using a microdrill (Stryker, Kalamazoo MI) and durotomy was performed to implant a removable silicone artificial dura precisely registered over motor cortex (Orsborn et al., 2015). A combination of machined ultem chamber parts (custom designs manufactured with Protolabs Inc.), silicone gaskets, and stainless steel machine screws were used to hermetically seal the chambers following surgery. Chambers were regularly inspected and cleaned in a sterile procedure room while the animals were awake and head-fixed in a primate research chair (Hybex Innovations, Montréal, CA). Prior to neural recordings, the artificial dura was swapped with one that contained embedded µECoG electrodes in a similar routine procedure.

### Electrophysiological and Behavioral Recordings

Neural signals were recorded using a 244-electrode µECoG array with 762 *µ*m inter-electrode pitch and 229 *µ*m contact size (Trumpis et al., 2021). The array was embedded in a silicone artificial dura to enable registration within the implanted chamber (Orsborn et al., 2015; Chiang et al., 2020). Once implanted, µECoG signals from 240 electrodes were amplified and digitized at 25 kHz using an eCube recording system (White-Matter Inc., Seattle WA).

Monkeys had head and left arm movements restricted as they made right arm reaches, which controlled the position of a computer cursor (Fig. 1A). The subject’s hand position was tracked in real-time using a marker-based camera system (NaturalPoint, Inc., Corvallis OR). The position of the hand was sent via multicast at 240 Hz to update the position of a cursor with which subjects performed goal-directed movements. The cursor was controlled by a custom version of BMI3D (http://github.com/aolabNeuro/brain-python-interface) and displayed on a 21.5 cm (measured vertically) 16:9 LCD display, which was updated at 120 Hz. The display was positioned 28 cm in front of the subject at eye level, directly above a 20 x 20 x 20 cm motion tracking volume in which monkeys could move their right arm. Coordinates within the tracking volume were mapped onto cursor position within a 20 x 20 cm square, centered on the display. Coordinates were scaled 1:1 so that hand movement up and to the right by 1 cm caused cursor movement up and to the right by 1 cm. Forward/backward movement of the hand relative to the screen had no impact on cursor movements. Display frame timing was output by BMI3D as a digital signal sent by microcontroller and recorded via the neural recording system to synchronize cursor and target timing with neural data. Display latency was measured with a Si photodiode (FDS100 from Thorlabs; mean 11, SD 4 ms after the digital sync).

**Figure 1.**
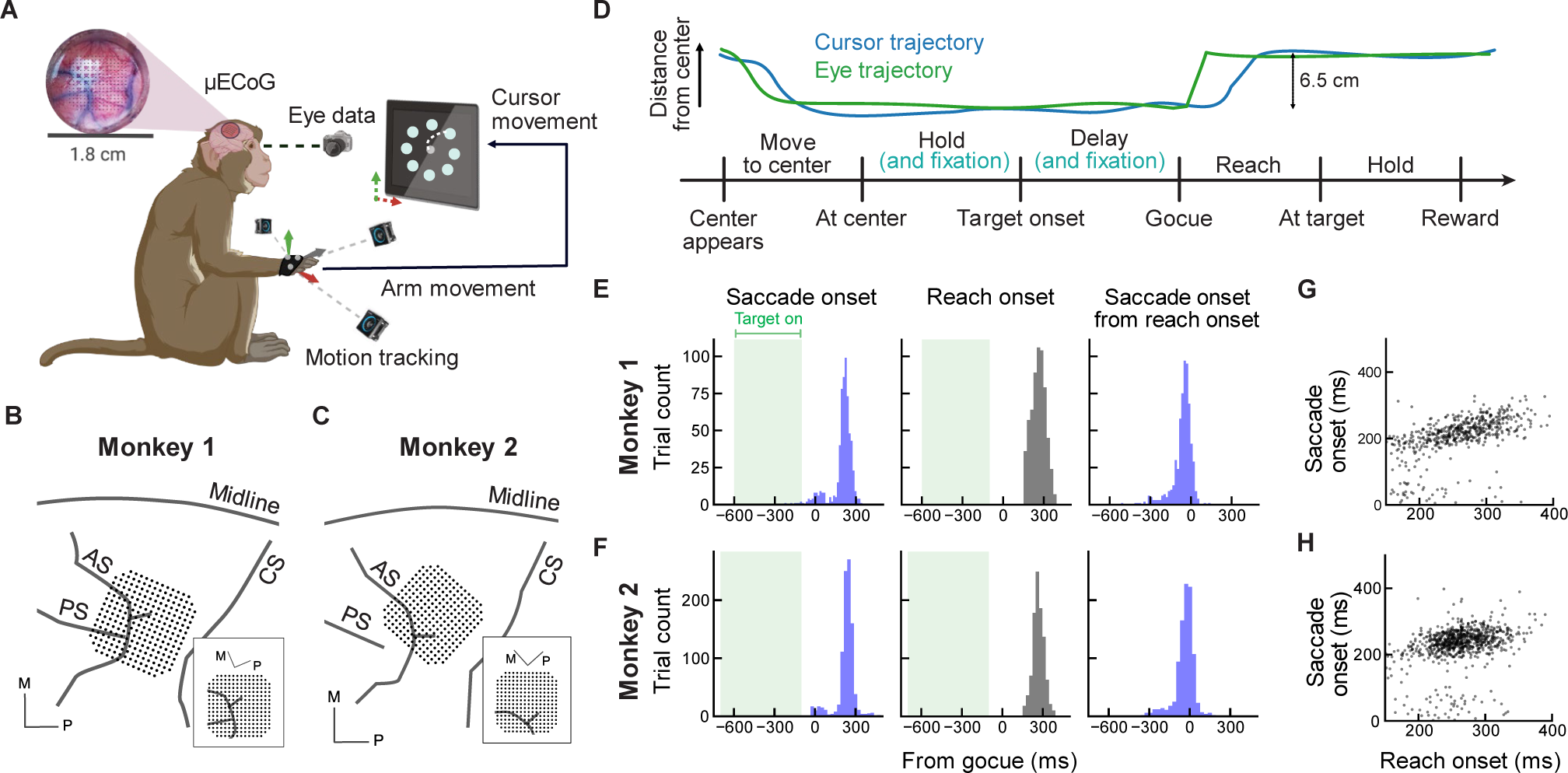
Experiment setup and behavioral data. **(A)** Schematic of the experimental setup. Arm movement was tracked and translated into the movement of a cursor on the visual display. **(B, C)** Schematic of recording locations of the µECoG array for Monkeys 1 and 2, respectively. Black lines indicate anatomical landmarks. AS: Arcuate sulcus, PS: Principal sulcus, CS: Central sulcus. M: medial, P: posterior. **(D)** Timeline of the delayed center-out reaching task (bottom) and an example of corresponding cursor and eye trajectories (top). Monkey 2 performed a modified version of this task which required fixating the center target during the initial hold period and delay period (cyan). **(E, F)** Histograms of saccade (left) and reach onset time (middle) following the go cue, and saccade onset time relative to reach onset time (right). Each monkey is shown independently in respective panels. Light green areas represent target onset times. Blue bars represent relevant saccade onset and black bars are movement onset. Shaded green regions show the range of target onset times. **(G, H)** The relationship between saccade onset times and reach onset times for Monkeys 1 and 2, respectively.

Images of subjects’ eyes were captured with an infrared camera at 240 Hz (FLIR, Wilsonville OR). Eye position within the camera frame was estimated in software (Zimmermann et al., 2016) and sent via digital-to-analog converters to the neural recording system. Eye position was subsequently calibrated to screen coordinates using regression between eye and cursor coordinates at the time of entering the peripheral targets. For Monkey 2, eye position within the camera frame was additionally sent via UDP packets to BMI3D for online gaze estimation.

Regression of the eye position to screen coordinates was computed on 100 successful center-out trials and subsequently used online to estimate the position of the animal’s gaze on the computer screen.

### Behavioral training and task

#### Initial behavioral training

Animals were trained in coordination with Behavioral Management Services at the Washington National Primate Research Center using positive reinforcement. Animals were selected from a pool based on availability and their willingness to interact with researchers. Following initial training, animals could sit quietly in a primate chair, have motion capture markers placed on their hands, and make reaching movements to a colored plastic ball held by a researcher.

#### Delayed center-out reaching task

Monkeys performed a self-paced delayed center-out reaching task (Fig. 1A and 1D). Task parameters used for each monkey can be found in Table 1. Trials began with the appearance of a center target. After a brief center hold, a peripheral target appeared 6.5 cm from the center target. Subjects were required to keep the cursor in the center target during a delay period, which was randomly drawn from a uniform distribution of the specific range. Disappearance of the center target signaled animals to acquire the peripheral target (the “go cue”). A successful trial required a brief hold at the peripheral target, after which a juice reward was delivered via silicone tubing routed inside flexible coolant line using a computer-controlled solenoid valve (custom design). The solenoid valve was grounded to the neural recording system to minimize electrical artifacts during rewards. Peripheral targets were presented in the pseudo-randomized order so that data for each target could be collected evenly.

**Table 1.** Task parameters for each monkey.

| Parameter | Monkey 1 | Monkey 2 |
| --- | --- | --- |
| Radius of target (cm) | 1.3 | 1.7 |
| Hold duration, center target (ms) | 200 | 150 |
| Range of delay period (ms) | 100 – 600 | 100 – 700 |
| Hold duration, peripheral target (ms) | 200 | 175 |
| Reward duration (ms) | 100 | 150 |

Training on the center-out task occurred over several hours per day and spanned 1.5 years (Monkey 1) or 0.75 years (Monkey 2), during which time animals were gradually introduced to smaller targets, longer delay periods, and finally smaller rewards until they were making more than 800 successful reaches in a single sitting. The animals’ eye positions were tracked during initial training, but they were not explicitly trained to fixate during any initial reach training.

After initial training in the delayed center-out reaching task, Monkey 2 exhibited a tendency to look away more frequently during the delay period compared to Monkey 1. We therefore modified Monkey 2’s task structure to require gaze fixation within a window centered around the center target (2.5 cm) until the go cue; eye movements outside of this fixation window caused the whole trial to restart. The color of the central target changed from yellow to cyan to indicate when the animal was successfully fixating. This modification resulted in more similar eye and arm behavior between Monkey 1 and 2 for cross-subject comparison. For Monkey 2, we recorded µECoG signals in variants of the task with and without fixation constraints in the delay period (controlled fixation condition (CF) and free fixation condition (FF), respectively).

Data from the CF condition was used for Monkey 2 in all analyses unless noted otherwise.

### Behavioral data analysis

Saccades were detected using a two-step thresholding procedure similar to prior studies (Tole and Young, 1981; Behrens et al., 2010). First, eye acceleration was estimated as the second derivative of the preprocessed eye position. Second, putative saccades were identified by detecting acceleration larger than a threshold acceleration. The threshold was determined for each trial as the mean plus 2.5 SD (0.7 SD for Monkey 2). Finally, to remove biologically implausible events, only putative saccades where acceleration increases and decreases occurred within a 15 - 160 ms window of each other were classified as saccades (Dorr et al., 2010).

Monkeys sometimes looked away during a trial, generating saccades that did not end with fixation to the peripheral target. Consequently, we used saccade start and endpoints to distinguish between task-relevant and task-irrelevant saccades. Saccades were deemed relevant if they originated within a 2.5 cm radius from the center target and ended within a 2.5 cm radius from the peripheral target. When multiple relevant saccades were detected, the saccade with the largest movement distance was defined as the relevant saccade. Any saccade other than the relevant saccade was classified as irrelevant.

Cursor movement onset (reach onset) after go cue was computed based on speed. The cursor speed was estimated by the first derivative of the preprocessed cursor positions, and digitally low pass filtered at 20 Hz. The times at which the filtered cursor speed exceeded a threshold speed (5 cm/s) were regarded as the reach onset.

### Neural data analysis

#### Preprocessing

Broadband neural data were filtered below 500 Hz using a fourth-order zero-phase Butterworth filter, then downsampled to 1,000 Hz. Data were then filtered from 0.1 Hz to 200 Hz using a 20,000th-order finite impulse response (FIR) filter to remove direct current components. Cursor position, recorded by BMI3D on each frame in screen coordinates, was interpolated to 25 kHz using the timing from the display. Cursor and eye kinematics were then decimated below 15 Hz and 200 Hz, respectively, using fourth-order zero-phase Butterworth filters, and resampled to 1,000 Hz.

#### Time-frequency analysis

We used a multi-taper method to calculate frequency band power from the µECoG signals. The preprocessed data were re-referenced using a common average across all channels (excluding those with poor signal quality). Spectral powers were computed between 0 Hz and 150 Hz in 10 ms steps using a 500 ms moving window, 4 Hz frequency band width, and 3 Slepian tapers. The time axis for spectral power was defined as the right edge of the time window in raw data. For example, raw data from the window spanning 0 ms to 500 ms generated the spectral estimate for time 0 ms in the spectral data. These multi-taper parameters were used in all analyses except for the inter trial-phase clustering analysis (see below), where a 100 ms window and 20 Hz frequency band width were used to increase temporal resolution. We averaged the spectral powers over each frequency range: delta (0.1-4 Hz), theta (4-8 Hz), alpha (8-14 Hz), beta (14-30 Hz), gamma (30-80 Hz) and high-gamma (80-150 Hz). These frequency ranges were determined based on the previous literature (Rickert et al., 2005; Schalk et al., 2007) and their approximate correspondence with frequency bands observed in our data (Fig. 2A and 2B).

**Figure 2.**
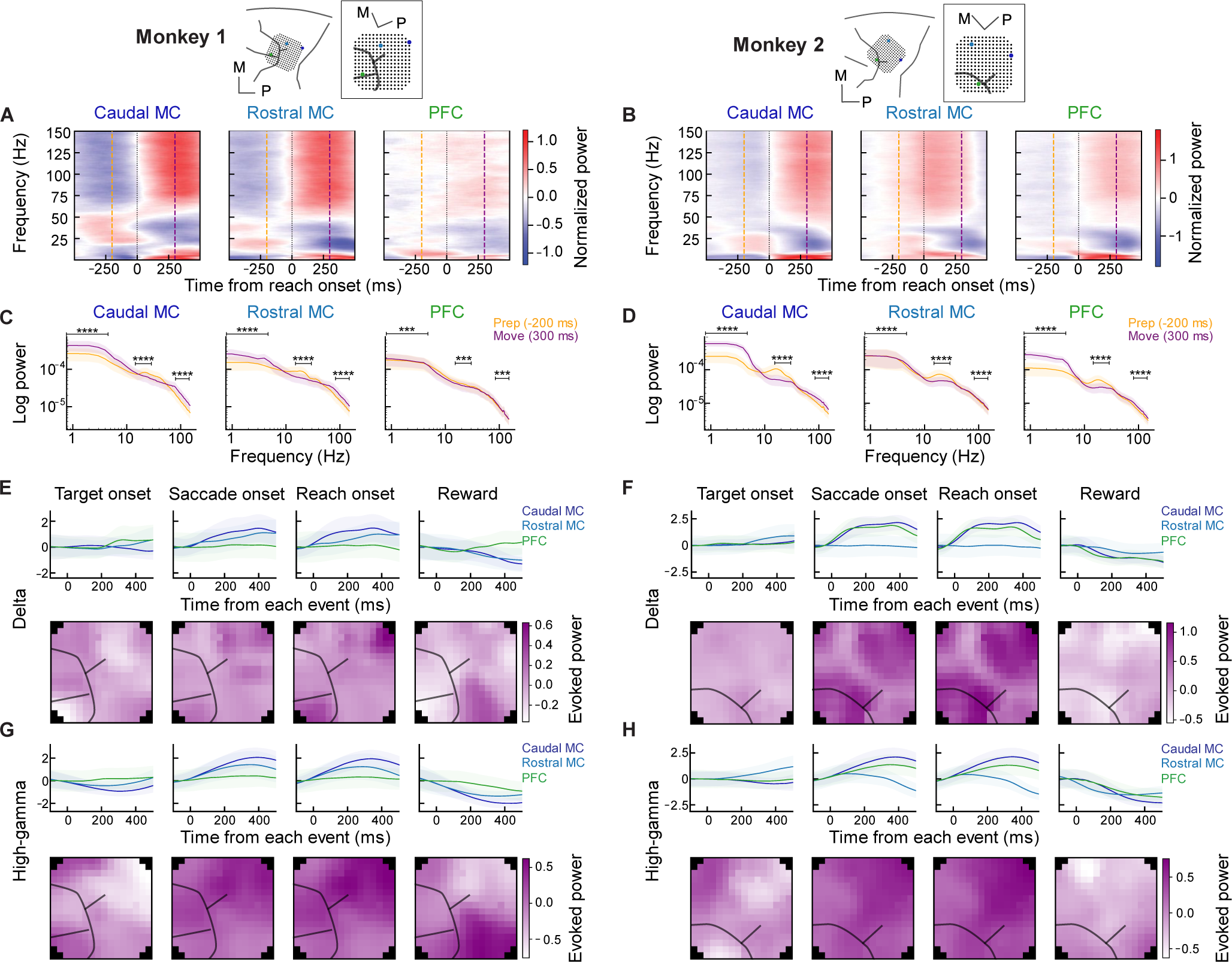
Time-frequency responses across brain areas show spatiotemporal patterns locked to multiple task features. **(A, B)** Mean spectrogram aligned to movement onset in example channels located in Caudal MC, Rostral MC, and PFC. Locations of channels are indicated in the top inset diagrams. Dashed lines indicate 200 ms before movement onset (orange, preparatory time) and 300 ms after movement onset (purple, movement time). **(C, D)** Log-scaled power spectra at 200 ms before (orange, preparatory time) and 300 ms after (purple, movement time) movement onset. Solid line indicates the mean and shaded area indicates the standard deviation across trials. **(E, F)** (Top) Evoked response in the delta band (0.1-4 Hz) power as a function of time for example electrodes around target onset, saccade onset, reach onset, and reward. Solid lines indicate the mean; shading represents the standard deviation across trials. (Bottom) Corresponding spatial map of the delta band power evoked response. Black lines indicate sulci. **(G, H)** Same as (E,F) but for the high-gamma band (80-150 Hz). For all sets of panels, each monkey is shown independently as indicated in the figure and the same example channels are illustrated throughout.

We also computed responses evoked by different task events in delta and high-gamma powers. First, delta and high-gamma powers were aligned to target onset, saccade onset, reach onset, and reward times. Data aligned to different task relevant events were then z-scored by using the mean and standard deviation for each channel separately, and these z-scored band powers were averaged across all trials. We calculated evoked powers by subtracting the mean power in a baseline period from the trial-averaged power. The baseline period was defined as [−100, 0] ms around each task relevant event. We also averaged evoked powers within an average time window (saccade onset, reach onset, and reward: [0, 100] ms; target onset: [150, 250] ms) to show spatial maps of evoked delta and high-gamma powers (Fig. 2E – 2H).

#### Direction tuning analysis

We analyzed information about movement target information within the delta power from 1.5 s before to 1.0 s after detected reach onset. We calculated a single mean and standard deviation across all data for each channel, which we used to z-score the delta power in each trial. Modulation depth (MD) was defined as the difference between maximum and minimum normalized power across target directions and was computed at every time point using a 100 ms moving window in 20 ms steps to examine how MD changed over time. Preparatory MD was computed within a preparatory interval ([−100, 0] ms around reach onset). Movement MD was computed within a movement interval ([200, 300] ms around reach onset). To identify the MD response time, we first defined a baseline period as [−1000, −750] ms before reach onset. We then calculated the time at which MD reached 7.0 standard deviations above the mean activity during the baseline period.

We employed the resampling method to calculate a mean for MD and response time. We generated a new sample by resampling trials without replacement and calculating the new mean MD and response time within the random sample. We repeated this procedure 300 times to create a resampling distribution. The mean and the standard deviation were obtained from this distribution.

#### Inter-trial phase clustering analysis

We computed inter-trial phase clustering (ITPC) to investigate inter-trial coherence in high-gamma power (Fig. 4A, Cohen, 2014). First, we developed a signal processing pipeline to extract reliable phase information from high-gamma signals. We computed the trial-averaged high-gamma power for all electrodes, and then calculated the average power spectral density (PSD) across channels to analyze the frequency content of high-gamma power. The trial-averaged high-gamma power primarily varied within a low frequency range (< 4 Hz, see Fig. 4A for an example). We therefore de-noised the high-gamma power by applying a band-pass filter from 0.5 to 4 Hz using a 1000^th^-order FIR filter. Subsequently, the Hilbert transform was applied to the filtered high-gamma power to extract phase information. The phase signals from each trial were extracted and aligned to saccade onset, reach onset, target acquisition, and reward. For each of these alignments we calculated ITPC at every time point:

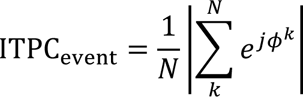

where ITPC_event_ is the ITPC aligned to that event (i.e. saccade onset, reach onset, target acquisition, or reward), *N* is the number of trials, *φ_k_* is a trial-aligned phase of the high-gamma activity at the *k*-th trial, and *j* is the imaginary unit. The ITPC ranges between 0 and 1.

If high-gamma power on a given channel was more related to eye movement than arm movement, phases of the high-gamma power should be more locked at saccade onset than reach onset. We calculated the effector preference index (EPI) between eye-aligned ITPC and arm-aligned ITPC to quantify how much eye-aligned ITPC differed from arm-aligned ITPC:

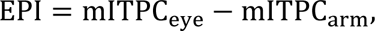

where mITPC_event_ is the event-aligned ITPC averaged within the analysis window ([−50, 50] ms). The EPI ranges between −1 and 1. Positive EPI indicates that the high-gamma phase of the channel was more consistently clustered around eye movement initiation than around arm movement initiation.

We similarly computed the reward index (RI) to quantify the relative change in trial consistency from target acquisition to reward events, as expressed by the following equation:

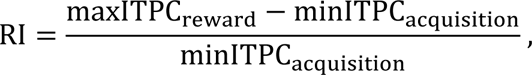

where minITPC_acquisition_ is the minimum ITPC_acquisition_ during the hold period and maxITPC_reward_ is the maximum ITPC_reward_ in the reward interval, which was defined as the period from the initiation of the reward until 300 ms after its completion. We computed the maximum or minimum of ITPC for RI instead of averaging it because ITPC continued to increase after reward times, making it difficult to define a time window for averaging. We also calculated non normalized RI as follows:

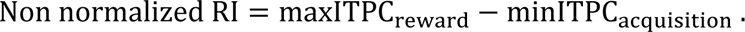

As with MD and response time, we employed the resampling method to calculate a mean for the ITPC magnitude at each timepoint, EPI and RI. We used the mean of the resampling distribution as the representative EPI and RI values. The 95% confidence interval (CI) was also obtained from the EPI and RI distributions.

#### Decoding analysis

We used linear discriminant analysis (LDA) to classify six spectral powers (delta: 0.1-4 Hz, theta: 4-8 Hz, alpha: 8-14 Hz, beta: 14-30 Hz, gamma: 30-80 Hz, and high-gamma: 80-150 Hz) across eight peripheral targets (Fig. 3I and Fig. 4H). First, we visualized how task information evolved over time by using neural activity from all channels, or channels selected based on anatomical location, to predict reach direction. We systematically varied the time window of neural data used to decode from 600 ms before to 600 ms after reach onset in 24.5 ms increments. Data was z-scored by using the mean and standard deviation across all data for each single channel and frequency band, ensuring equal contribution of frequency power from each channel to the decoding analysis. To decode target directions at a given time point, the z-scored data was averaged from [−100, 0] ms relative to that time point, then input into LDA. LDA was implemented using the Python library Scikit-learn (Pedregosa et al., 2011) with the eigen solver and automatic shrinkage.

**Figure 3.**
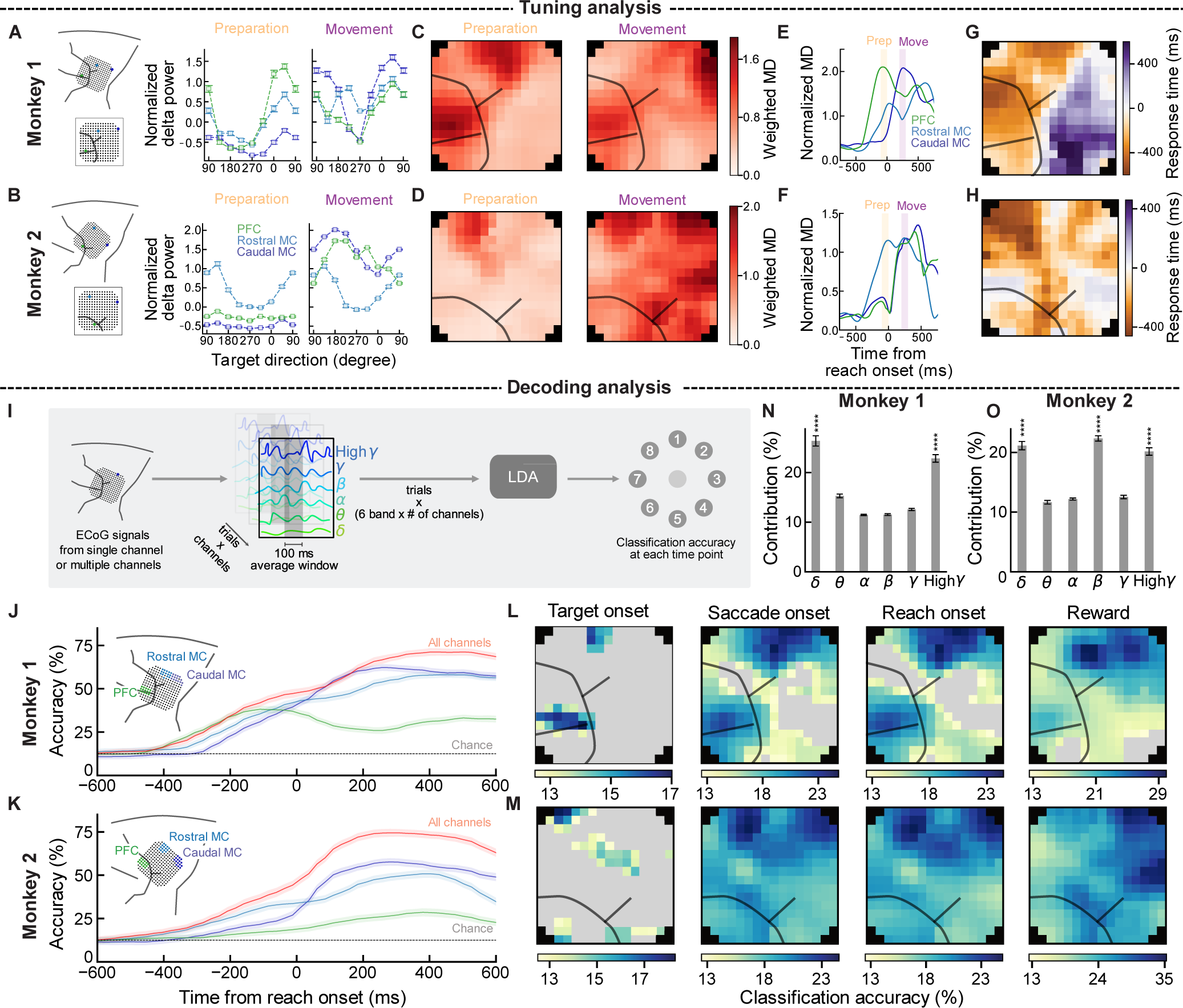
Directional information shifts from rostral to caudal in frontal cortices following arm movement onset. **(A, B)** Mean normalized delta power as a function of movement target direction before reach onset ([−100, 0] ms) and after reach onset ([200, 300] ms) in example channels in Caudal MC (dark blue), Rostral MC (blue), and PFC (green). Insets on the left show the array position with anatomical landmarks and indicate the location of example electrodes. **(C, D)** Spatial map of modulation depth before (left) and after (right) reach onset. Black lines indicate approximate sulci locations. **(E, F)** Modulation depth as a function of time for example channels in Caudal MC (dark blue), Rostral MC (blue), and PFC (green). **(G, H)** Spatial map of response time calculated based on modulation depth. Black lines indicate approximate sulci locations. **(I)** Schematic of decoding analysis. Features extracted from single or multiple ECoG channels were used as input to a LDA model to determine classification accuracy at each time point. **(J, K)** Classification accuracies at each time point when data was aligned to reach onset for all channels (red), Caudal MC (dark blue), Rostral MC (blue), and PFC (green). Inset at the left show the channels used for the decoding analysis. Solid line indicates the mean and shading indicates the standard deviation across the resampling distribution. **(L, M)** Spatial maps of single channel classification accuracy at different task epochs: target onset, saccade onset, reach onset and reward. Gray channels represent channels with statistically insignificant classification accuracy. **(N, O)** Contribution of each frequency band to classification accuracy based on LDA model weights. Bar height indicates the mean across channels, error bars indicate s.e.m. across channels. For all sets of panels (excluding I), each monkey is shown independently as indicated in the figure.

We analyzed the spatial distribution of task information by calculating target decoding accuracy from single µECoG channels. Decoding was performed using the same six frequency band powers, z-scored across all data for each channel and frequency band, recentered to task-events (target onset, saccade onset, reach onset, and reward times) and averaged over task-relevant time windows (saccade onset, reach onset, and reward: [−50, 50] ms, target onset: [150, 250] ms).

We estimated decoding accuracy using the resampling method and 5-fold cross-validation. Decoding accuracy (ratio of correctly labeled trials to the total number of trials) in the test set was calculated as the average across the 5 folds. We repeated this sampling procedure 300 times to create a resampling distribution of decoding accuracy. All reported results reflect the decoding accuracy averaged across the resampling distribution. Error bars were calculated as the standard deviation of the distribution (Fig. 3J and 3K).

To compute significance of decoding accuracies at each channel, we used permutation tests in which target labels were randomly shuffled to estimate the chance (N = 300). We then calculated the statistical significance of decoding accuracy (*p* values) by determining the probability of obtaining the mean of decoding accuracy from the null distribution (Fig. 3L and 3M).

We assessed the contribution of each frequency band to target decoding by analyzing the LDA model coefficients (Fig. 3N and 3O). For this analysis we used the decoding accuracies that were calculated on data aligned to reach onset ([−50, 50] ms window). Decoding contribution was calculated by averaging the model coefficient weights across resampled samples, folds, and channels for each frequency band.

We also investigated how gaze behavior influenced µECoG signals by comparing task decoding performance between two separate sessions recorded in Monkey 2 (Fig. 4H). In the first session, eye positions were controlled during the delay period (CF), while the second session allowed for free fixation (FF). We used the resampling method to control for differences in the total number of trials between session types. We randomly drew the same number of trials from each session without replacement (N=300). Target direction was predicted from 6 frequency band powers averaged within preparatory times ([−200, 0] ms around reach onset) and movement times ([200, 400] ms around reach onset). The resulting data was fed into the LDA and decoding accuracies in the two sessions were estimated using 5-fold cross-validation. We also tested whether there were statistically significant differences between the two decoding accuracies from CF and FF. The distribution of differences in the decoding accuracy was calculated through a resampling procedure. These distributions were used to estimate *p*-values to reject the hypothesis that there is no statistically significant difference. Specifically, if the mean of the differences was greater than 0, we calculated the proportion of samples in the resampled distribution that were less than 0. (When the mean was less than zero, we calculated the proportion of samples in the resampled distribution that were greater than 0.) This proportion of samples was then doubled to make it comparable to a two-sided test.

**Figure 4.**
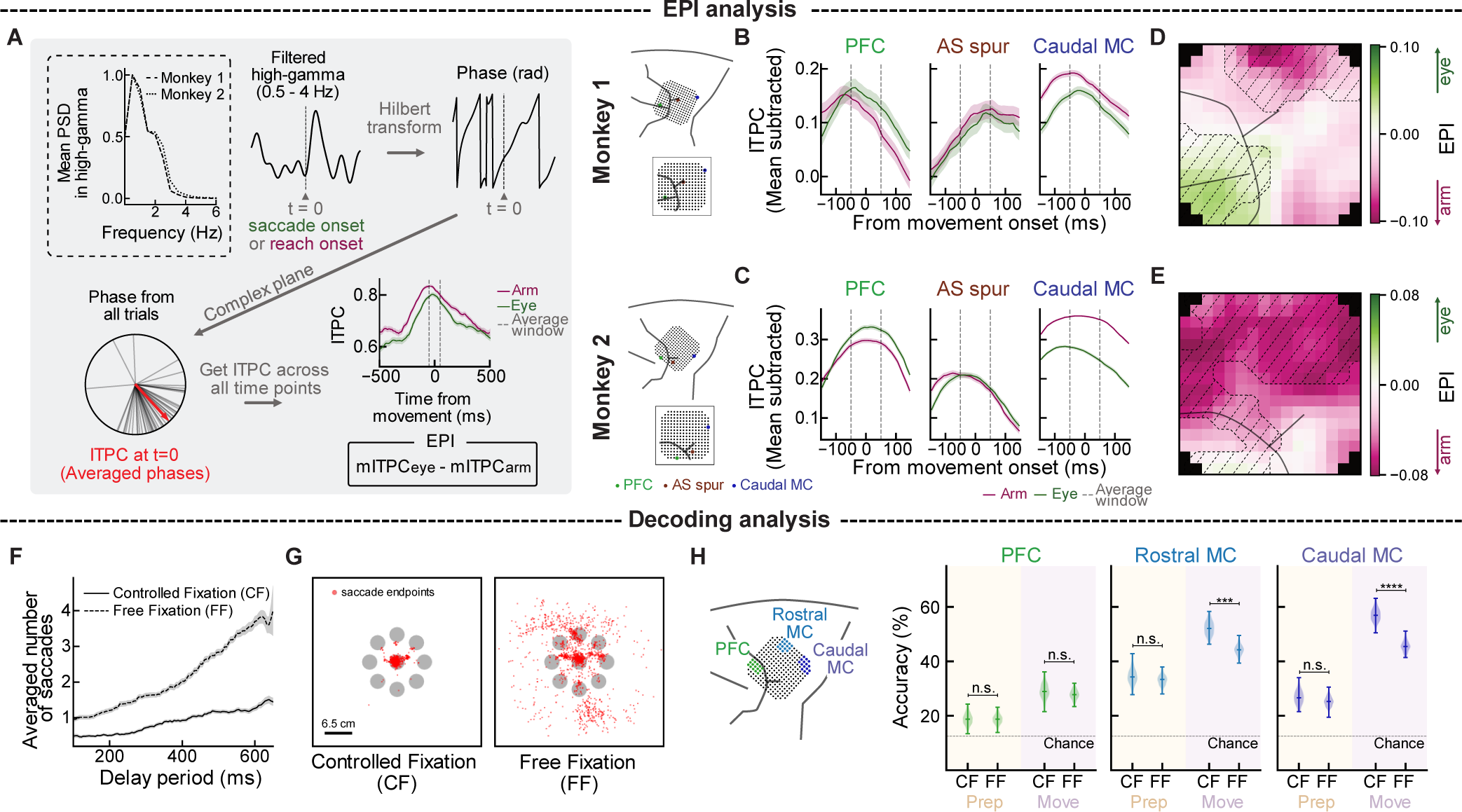
Overlap of eye and arm information in the motor cortex. **(A)** Schematic of the Effector Preference Index (EPI) analysis with data from an example channel for each calculation step. Mean PSD in trial-averaged high-gamma power was computed to determine the frequency band for band-pass filtering (top left). After high-gamma power was band-pass filtered between 0.5 – 4 Hz, the Hilbert transform was applied to extract phase information in each trial. Phases were averaged across all trials on the complex plane to compute ITPC. ITPC was calculated for eye- and arm-aligned high-gamma power. Finally, EPI was computed by comparing eye-aligned ITPC and arm-aligned ITPC around 0. **(B, C)** Eye-aligned ITPC (green) and arm-aligned ITPC (pink) for example channels located in PFC, near AS, and within Caudal MC for monkeys 1 and 2, respectively. Solid line indicates the mean and shading indicates the standard deviation estimated from resampling distributions. Black dashed lines indicate the average time window to compute EPI. Insets on the left show the array position with anatomical landmarks and indicate the location of example electrodes. **(D, E)** Spatial map of EPI for monkeys 1 and 2, respectively. Squares with different colors indicate the location of the three channels shown in (B, C). Black solid lines represent approximate sulci locations. Black dashed lines show high decoding accuracy areas around reach onset more than 16% for Monkey 1 and 18% for Monkey 2 from Fig. 3. **(F)** Average number of saccades as a function of the trial delay period length for Monkey 2 in the controlled fixation (CF, solid black line) and free fixation (FF, dashed black line) task conditions. Solid line indicates the mean; shaded region indicates s.e.m. over a 100 ms moving window. **(G)** End points of saccades for select trials during controlled (left) and free fixation (right) task (monkey 2). **(H)** Target classification accuracy using channels from PFC (green), rostral MC (blue) and Caudal MC (dark blue) during preparatory and movement periods in controlled vs free fixation. Error bars indicate confidence interval estimated using 300 bootstraps. Inset at the left depict the spatial locations of channels used for decoding analysis from each region.

### Data selection

The present analysis included data from a single day (Monkey 1), two days (Monkey 2 controlled fixation), or five days (Monkey 2 free fixation). Behavioral sessions were selected to maximize neural signal quality and the number of successful reaches.

To account for the possibility that monkeys anticipated the go cue or missed its appearance, trials with movement onsets less than 150 ms or more than 400 ms after the go cue were excluded from the dataset (Churchland et al., 2006; Batista et al., 2007). Additionally, trials lacking any relevant saccades were excluded to minimize the influence of inconsistent eye behavior on neural data. This selection procedure identified 533 of the 800 successful trials initially collected for Monkey 1 and 829 out of 1078 in Monkey 2. For the decoding analysis shown in Fig. 4H we included trials without any relevant saccades to examine how irrelevant eye movement affected decoding performance (1017 success trials from 1078 success trials in CF condition, 507 success trials from 792 success trials in FF condition).

µECoG recording electrodes with poor signal quality were identified and removed. First, the variability of preprocessed neural data was calculated for each channel using the standard deviation of the first 60 seconds of data. Channels were kept if their variability fell within the variability range of 5% to 95% of all channels. Second, the remaining channels were classified as having poor signal quality if their variability exceeded 5 times the median variability across all remaining channels. We kept 238 out of 244 channels for Monkey 1, 233 out of 244 channels in both CF and FF conditions for Monkey 2.

### Statistical analyses

We recorded data and presented results from 2 monkeys. Results were not pooled across the two monkeys. Data were collected with a consistent number of successful trials per target. This uniformity collapsed when we extracted trials based on trial selection criteria (See Data selection). We used all trials that satisfied trial selection criteria only for Fig. 2 because spectrogram and evoked powers in Fig. 2 were averaged across all trials and the uniformity for target directions was not necessary. However, when we investigate neural activity that depends on target directions such as tuning curve analysis, uniformity was required. Opting for arbitrary trials to maintain uniformity just once could result in a biased conclusion. We therefore employed the resampling method for all other analyses. After the trial selection based on behavioral data, the minimum number of trials per target was 51 trials for Monkey 1 and 92 trials for Monkey 2. Therefore, we chose the minimum number of trials per target for each target condition without replacement so that the number of trials for each condition would be the same across conditions. For decoding analysis in CF and FF sessions for Monkey 2, 54 trials for each target condition were chosen without replacement to make the number of training samples the same across sessions. The resampling procedure was repeated 300 times. Details of the resampling method and the statistical tests we conducted for each analysis are provided in the respective sections of Behavioral data analysis and Neural data analysis. Significance levels are indicated as follows: **** : p≤10^−4^, *** : 10^−4^<p≤10^−3^, ** : 10^−3^<p≤10^−2^,* : 10^−2^<p≤0.05, ns : p>0.05.

### Visualization of spatial maps

Spatial maps of modulation depth and decoding accuracy were visualized by scaling data based on statistical significance. The weighting mask, denoted as w, was calculated as:

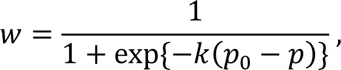

where *p* denotes the estimated *p*-value of the data, and *k* and *p_0_* are constants set to 100 and 0.08, respectively (Chiang et al., 2020). This defines a weight *w* that was close to 1 for small *p*-values but quickly decayed to 0 as the *p*-values increased, effectively masking statistically insignificant “noise”. To aide visualization, missing or excluded data were interpolated based on values from neighboring channels for all spatial maps. Finally, maps were smoothed by averaging across neighboring channels (up to 9 channels). All scaling and smoothing was applied after statistical tests were performed. For Fig. 3L and 3M, the decoding accuracy at single channels was set to NaN when the scaled decoding accuracy became below 100/8 (chance level), which are shown as gray color.

## Results

We trained two male macaque monkeys to perform a self-paced delayed center-out reaching task (Fig. 1A and 1B, see Materials and Methods) to investigate simultaneous eye movement, movement preparation, movement execution, and receipt of rewards. We recorded neural activity using a 244-channel ECoG array positioned sub-durally over the prefrontal cortex and motor cortex (Fig. 1C). These areas were chosen because they are involved in eye movement, arm movement preparation, and arm movement execution (Georgopoulos et al., 1982; Crammond and Kalaska, 2000; Schall, 2002; Funahashi, 2014).

Since eye and arm movements are often coordinated, we examined the temporal relationship between saccade onset and arm movement onset (Fig. 1D). Saccades were classified as either relevant or irrelevant to the task (See Materials and Methods). Trials without relevant saccades were excluded. Both monkeys showed relevant saccades, although Monkey 2 generated more irrelevant saccades even in the controlled fixation condition (ratio of relevant to irrelevant saccades: 704/907 for Monkey 1, 882/4387 for Monkey 2). Relevant saccade onset happened after the go cue (median ± SD: 220 ms ± 114 ms for Monkey 1, 247 ms ± 169 ms for Monkey 2; Fig. 1E, left). Reach onset also occurred after the go cue (median ± SD: 262 ms ± 52.3 ms for Monkey 1, 265 ms ± 41.7 m for Monkey 2; Fig. 1E, middle). Eye movements were initiated earlier than arm movements (median difference between saccade and reach onsets ± SD: 47.0 ms ± 106 ms, 13.0 ms ± 161 ms, Wilcoxon signed-rank test: p = 5.49 × 10^-75^, p = 8.03 × 10^-52^, for Monkey 1 and Monkey 2, respectively; Fig. 1E, right). Saccade sometimes happened before the go cue. We excluded these trials (29 and 24 trials for Monkeys 1 and 2, respectively) to examine the correlation between reaction times for saccades and reaches. Saccade onset times were correlated with reach onset times (Pearson correlation coefficient: 0.520, p = 1.63 × 10^-37^ for Monkey 1, 0.287, p = 1.00 × 10^-23^ for Monkey 2; Fig. 1G and 1H). This correlated movement structure was preserved even after we included trials where saccade occurred earlier than go-cue (Pearson correlation coefficient: 0.469, p = 1.54 × 10^-31^ for Monkey 1, 0.186, p = 7.30 × 10^-8^ for Monkey 2). These results show that the timing of arm movement and relevant saccades followed a consistent and commonly-observed structure where the monkeys saccade to the peripheral target shortly before initiating an arm movement as part of a coordinated movement plan (Dean et al., 2011).

### Spatial variations in neural responses across brain regions

Next, we investigated the spectral features of neural activity captured by our µECoG array across motor cortex (MC) and prefrontal cortex (PFC), which have been shown to relate to eye and arm movements (Lee et al., 2017; Volkova et al., 2019). We first characterized the general frequency structure by calculating average spectrograms across all selected trials, aligned to reach onset (Fig. 2A). Example channels from different regions (Caudal MC, Rostral MC, PFC) were chosen based on anatomical locations. We observed three visually distinct frequency bands in both animals. For instance, in Caudal MC of Monkey 1, there were distinct bands of activity in 0-20 Hz, 20-50 Hz, and 50-150 Hz bands (Fig. 2A). The frequency ranges approximately correspond to commonly studied frequency bands: delta (0.1-4 Hz), beta (14-30 Hz), and high-gamma (80-150 Hz). We therefore used the frequency ranges defined in previous literature for subsequent band-specific analyses (Rickert et al., 2005; Schalk et al., 2007). We also visualized movement-related differences in spectral power by comparing the log-scaled spectral power during preparation (200 ms before go cue) versus during arm movement (300 ms after go cue) (Fig. 2C and 2D). We observed changes in power across all frequency bands of interest (delta, beta, and high-gamma) in many electrodes (Quantified for 3 example electrodes for each monkey; Wilcoxon signed-rank test, Monkey 1: p<1×10^-3^, Monkey 2: p<1×10^-4^ for all frequency bands). These data are consistent with prior reports (Miller et al., 2007; Ball et al., 2009).

Previous literature showed that delta and high-gamma band power contain movement information (Rickert et al., 2005). Our results also showed statistically significant changes before and after reach onset in delta and high-gamma powers. We first examined how these spectral features relate to task relevant events: target onset, saccade onset, reach onset, and receipt of reward (Fig. 2E – 2H). Delta band power showed evoked responses for all task events (Fig. 2E and 2F, top) with notable spatial variability across brain regions (Fig. 2E and 2F, bottom). Consistent with the tight temporal relationship between saccade and reach onset, evoked responses and their spatial distributions for these events were qualitatively similar.

High-gamma power also showed evoked responses for all task events (Fig. 2G and 2H, top) that spatially varied across brain regions (Fig. 2G and 2H, bottom) and were qualitatively similar for saccade and reach onset. These analyses show that µECoG signals across PFC and MC contain spatially distributed information about a mixture of computations, from visual responses to targets, movements, and rewards. However, these spectral analyses alone are also limited in their ability to separate information about each computation.

### Mapping the temporal evolution of movement direction information

Having established temporal relationships between saccades and arm movements (Fig. 1) and the spatiotemporal patterns of gross task information (Fig. 2), we next explored how information about the movement location was distributed across brain areas over time. We first analyzed movement direction information by computing tuning curves (see Methods) across electrodes and at different times within the task. We first focus on signals in the delta band (0.1 - 4 Hz) based on our observations above (Fig. 2E, 2F) and past reports of its’ utility for direction decoding (Rickert et al., 2005; Chiang et al., 2020).

We compared tuning curves before and after reach onset ([−100, 0] ms and [200, 300] ms, respectively). We then computed the modulation depth (MD) for each electrode to quantify the spatial distribution of directional tuning. Example tuning curves are shown in Fig. 3A and 3B. Prior to reach onset, tuning was larger in rostral MC compared to caudal MC, consistent with rostral MC’s role in motor planning (Table 2). Tuning in the PFC prior to reach onset varied across monkeys. After reach onset, three example channels (PFC, Rostral MC, and Caudal MC) showed clear direction tuning (Table 2). These trends of information moving from rostral to caudal MC and variable directional information in PFC across monkeys were also clear from comparing MD maps before and after reach onset (Fig. 3C, 3D). These analyses suggest that µECoG maps can capture shifts in movement-related processing across the frontal motor cortices as the animal shifts from planning to executing arm movements. We quantified shifts in movement information across brain regions over time by calculating MD at every time point for each electrode (see Materials & Methods). Example electrodes illustrate differences in the timing of directional information across brain regions (Fig. 3E, 3F). A spatial map in the timing of directional tuning revealed spatial structure across the cortical surface (Fig. 3G, 3H). In both monkeys, tuning emerged earlier in rostral MC compared to caudal MC. Interestingly, while the onset of directional tuning in PFC was similar for both monkeys, the timing of the largest directional tuning in PFC varied between animals (Fig. 3E, 3F). In Monkey 1, PFC was tuned prior to reach onset, while in Monkey 2, PFC was tuned slightly before reach onset and its large tuning was more closely tied to reach onset (along with caudal MC) (Fig. 3E, 3F).

**Table 2.** Modulation depth (MD) at preparatory and movement times across different channels.

| Monkey | Channel | Median of MD at preparatory times | Median of MD at movement times |
| --- | --- | --- | --- |
| 1 | Rostral MC | 1.25 | 1.04 |
|  | Caudal MC | 0.618 | 2.05 |
|  | PFC | 2.04 | 1.44 |
| 2 | Rostral MC | 1.15 | 1.14 |
|  | Caudal MC | 0.242 | 1.17 |
|  | PFC | 0.244 | 1.12 |

While these band-specific analyses are powerful, they may not fully capture all task information represented in µECoG signals. We next used linear discriminant analysis (LDA) decoding to explore how information about the movement target was represented across brain areas and frequency bands. We grouped channels by anatomical location (PFC, Rostral MC, caudal MC, and all electrodes) and then used 6 different frequency bands to decode 8 target directions (Fig. 3I). In both monkeys, using all channels led to high decoding accuracy shortly after reach onset (maximum decoding accuracy: Monkey 1, 69.9%; Monkey 2, 72.8%, Fig. 3J, 3K). In both monkeys, the decoding accuracy of Rostral MC was higher than Caudal MC before arm movement onset. The PFC also had directional information for Monkey 1, but less information for Monkey 2 in the delay period. The decoding accuracy increased after reach initiation across all brain regions.

We then calculated single channel decoding accuracy to obtain high-resolution spatial maps of where target information is represented (Fig. 3L, 3M). Comparing the decoding accuracy maps across different phases of the task (target onset, saccade onset, reach onset, and reward) revealed how task information propagates across brain areas. In both monkeys, channels in PFC and Rostral MC had statistically significant classification accuracy around target onset.

The number of channels with high decoding accuracy within Rostral MC increased at saccade onset. At reach onset, directional information was mainly located in Rostral MC and Caudal MC. Target information stayed in Caudal MC around the time of reward (Fig 3L, 3M). The mean single channel decoding accuracy also increased as the task phase progressed (mean ± SD at target onset, saccade onset, reach onset and reward: 13.8±1.46%, 17.7±3.26%, 17.5±3.23%, 19.2±4.46% for Monkey 1; 13.9±1.33%, 19.4±2.30%, 19.5±2.62%, 25.5±4.78% for Monkey 2). We also found differences across monkeys. For Monkey 1, many channels in PFC had high directional information from target onset to reach onset. Rostral MC also had information even in reward times (Fig. 3L). For Monkey 2, movement related information was more distributed across all electrodes at saccade onset, reach onset, and reward (Fig. 3M, the number of significance channels at target onset, saccade onset, reach onset, and reward: 42, 196, 194, 220 out of 238 channels for Monkey1; 93, 232, 232, 233 out of 233 channels for Monkey 2). PFC had less information compared to Rostral MC and Caudal MC in reward times (Fig. 3M).

Lastly, analyzing the classifier weights allowed us to examine which frequency band(s) encoded target directions (Fig. 3N, 3O). The delta and high-gamma band had statistically significant directional information (Monkey1: *p* = 1.14×10^-24^ and *p* = 3.96×10^-34^ for delta and high-gamma, Monkey 2: *p* = 5.21×10^-12^ and *p* = 1.96×10^-12^, for delta and high-gamma, One-sided Wilcoxon signed-rank test). This result supports movement related information within delta and high-gamma bands. Notably, Monkey 2 had high beta band contribution (Monkey1: *p* =0.115, Monkey2: *p* = 3.43×10^-27^, Wilcoxon signed-rank one-sided test), which may reflect different task demands (see Discussion).

Together, our results show that information about movement direction is present in both PFC and MC, and that µECoG signals can map how this information shifts across the brain over time. Specifically, directional information shifts from Rostral MC and PFC to Caudal MC over the course of the reach. These results, paired with past work illustrating the roles of PFC and MC in eye and arm movements, suggest this target information likely reflects processing related to preparing and initiating both eye and arm movements.

### Eye and arm movement information overlaps in frontal cortices

We observed changes in directional information around reach onset in both PFC and MC (Fig. 3). This directional information could be related to preparing and executing eye movements, arm movements, or both. Understanding whether these variations are associated with eye movements or arm movements is essential to parse the neural computation underlying these movements. However, eye and arm movement behaviors were closely correlated (Fig. 1G, 1H). As a result, comparing evoked power (Fig. 2E-2H) or target information (Fig. 3L, 3M) aligned to saccade or reach onset was not able to distinguish between effectors.

To resolve differences in neural activity related to eye movements versus arm movements, we turned to inter-trial phase analyses that are more sensitive to timing differences. We used inter-trial phase clustering (ITPC) to quantify the consistency in the phase of high-gamma activity, trial-to-trial. If a spectral feature within the ECoG is more correlated with eye movement, the phase of that signal should be more consistent (i.e., clustered) around eye movement onset than arm movement initiation. This would result in ITPC aligned to eye movement onset (eye-aligned ITPC) that is higher than ITPC aligned to arm movement onset (arm-aligned ITPC). We defined an Effector Preference Index (EPI, schematized in Fig. 4A), which was calculated for each channel as the difference between the eye-aligned ITPC and the arm-aligned ITPC (See Materials and Methods) within the high-gamma power. If the phase of high-gamma power is more consistently aligned to eye movements, the EPI is a positive value; negative EPI values correspond to more consistent alignment to arm movements.

Examples of ITPC and EPI in three different brain areas are depicted in Fig. 4B, 4C and Table 3. In PFC, the eye-aligned ITPC exceeded the arm-aligned ITPC around movement onset. On the other hand, in Caudal MC, the eye-aligned ITPC was lower than arm-aligned ITPC around movement onset around the AS spur, the sign of EPI was not consistent across monkeys. However, the absolute value of EPI was smaller, and its 95% CI included 0. These examples suggest a spatial difference in which areas have information about each effector.

**Table 3.** EPI and CI in three representative channels.

| Monkey | Channel | EPI | 95% CI |
| --- | --- | --- | --- |
| 1 | PFC | 0.029 | [0.0056, 0.052] |
|  | AS spur | -0.015 | [-0.041, 0.0096] |
|  | Caudal MC | -0.027 | [-0.042, -0.012] |
| 2 | PFC | 0.032 | [0.022, 0.042] |
|  | AS spur | 0.0049 | [-0.0046, 0.014] |
|  | Caudal MC | -0.090 | [-0.098, -0.083] |

Fig. 4D and 4E show EPI maps with black shaded areas that represent high decoding accuracy channels around reach onset displayed in Fig 3L and 3M. The EPI maps revealed clear spatial structure in where eye and arm information were located across brain regions (Fig. 4D, 4E). Channels with a higher EPI were predominantly located in PFC. Premotor regions around the AS spur also exhibited a high EPI, but also had smaller absolute values of EPI than PFC and MC. Negative EPI channels, reflecting more significant arm representations, were found in portions of Rostral MC for both monkeys. These channels were more distributed across the motor area for Monkey 2. These maps show slight differences between monkeys, with Monkey 2 showing few electrodes with a strong eye preference compared to Monkey 1.

This may be related to the timing of eye and arm movements, which were slightly closer in Monkey 2 than Monkey 1 (median difference between saccade and reach onsets ± SD: 47.0 ms ± 106 ms, 13.0 ms ± 161 ms, Fig. 1E). These results show that high-gamma activity in PFC is more temporally linked to eye movements, while motor areas are more temporally linked to arm movement, and the premotor area near the AS was temporally related to eye movement or both effectors.

Integrating the EPI and target decoding analyses suggests that both eye and arm movement information contributed to target decoding. Channels with high decoding accuracy around reach onset were found in PFC, Rostral MC, and Caudal MC for both monkeys. EPI results suggest that directional information in PFC may predominantly reflect information about eye movements, while motor areas may be biased towards information about arm movements.

To more directly test whether eye and arm information were separated across these regions, we performed decoding analyses across different gaze conditions. If information about eye and arm movements is mixed within µECoG signals in frontal cortices, we would predict that gaze position will influence the decoding of arm movements. Indeed, past work showed that offline decoding accuracy of arm movements from neuronal spiking activity in pre-motor cortices was higher when gaze position was systematically controlled (Batista et al., 2008). We tested this prediction by comparing target decoding in sessions with and without fixation requirements during the delay period (controlled fixation condition – CF and free fixation condition – FF, respectively) in Monkey 2. We performed target decoding analysis using multiple channels for each session and quantified the difference in decoding performance between the two conditions across brain regions.

We first compared the monkey’s behavior in the two sessions. In the CF condition, the average number of saccades before go cue was consistently less than one, consistent with holding fixation during the delay period (Fig. 4F). In the FF condition, the monkey made more saccades in the delay period (mean absolute difference in the number of saccades between the two conditions, 1.27, *p* = 3.90×10^-18^, Wilcoxon signed-rank test). Next, we compared the saccade end positions in the delay period between the two conditions (Fig. 4G). In the controlled fixation condition, saccades were sometimes generated, but most of them remained within the center target. In the free fixation condition, the end positions were more scattered (differences in mean distance of saccade end positions, 5.67 cm, *p* = 4.56×10^-232^, Wilcoxon rank-sum test). These results show that the monkey frequently generated saccades to locations other than the peripheral target in the FF condition. We predict these eye movements to task-irrelevant locations will reduce our ability to decode the location of the peripheral target.

We quantified how task-irrelevant eye movements affected decoding performance by comparing decoding accuracy at preparatory ([−200, 0] ms around reach onset) and movement ([200, 400] ms around reach onset) periods between the two task conditions. These periods were chosen based on decoding accuracy of the CF data (Fig. 3K). We performed target decoding using the same groups of channels in Caudal MC, Rostral MC, and PFC as in Fig. 3K for both sessions. In Caudal MC and Rostral MC, the decoding accuracy was higher in the CF condition compared to the FF condition at movement times (PFC: *p* = 0.687, Rostral MC: *p* = 6.67×10^-3^, Caudal MC: *p* = 0.0, estimated from resampling distributions). Decoding accuracy during the preparatory period was not statistically different between the two conditions across all brain regions (PFC: *p* = 0.973, Rostral MC: *p* = 0.727, Caudal MC: *p* = 0.647, estimated from resampling distributions). These findings show that task-irrelevant eye movements impacted decoding performance within motor cortices, which indicates the presence of eye related activity across motor regions, including caudal areas near the primary motor cortex.

### Reward related information in the frontal cortex is localized

In addition to movement related activity, frontal cortices concurrently encode reward-related activity (Roesch and Olson, 2003; Ramkumar et al., 2016; Ramakrishnan et al., 2017), which likely contribute to shaping eye and arm movements during motor learning. While reward-related information has been observed in different frontal cortical areas, the spatial distribution is not well-characterized. We performed analyses to resolve differences in the temporal relationships between µECoG signals and reward to generate functional maps of reward related signals across PFC and MC.

We again used phase-related analyses to examine how neural activity related to reward (Fig. 5A). If the neural activity at a given channel is related to reward, then the extent of phase clustering in high-gamma power should be higher to around reward times than before the reward (during target acquisition). We used ITPC to quantify consistency of high-gamma power across trials. We defined a Reward Index (RI) as the relative magnitude of ITPC increase from target acquisition to reward times (See Materials and Methods).

**Figure 5.**
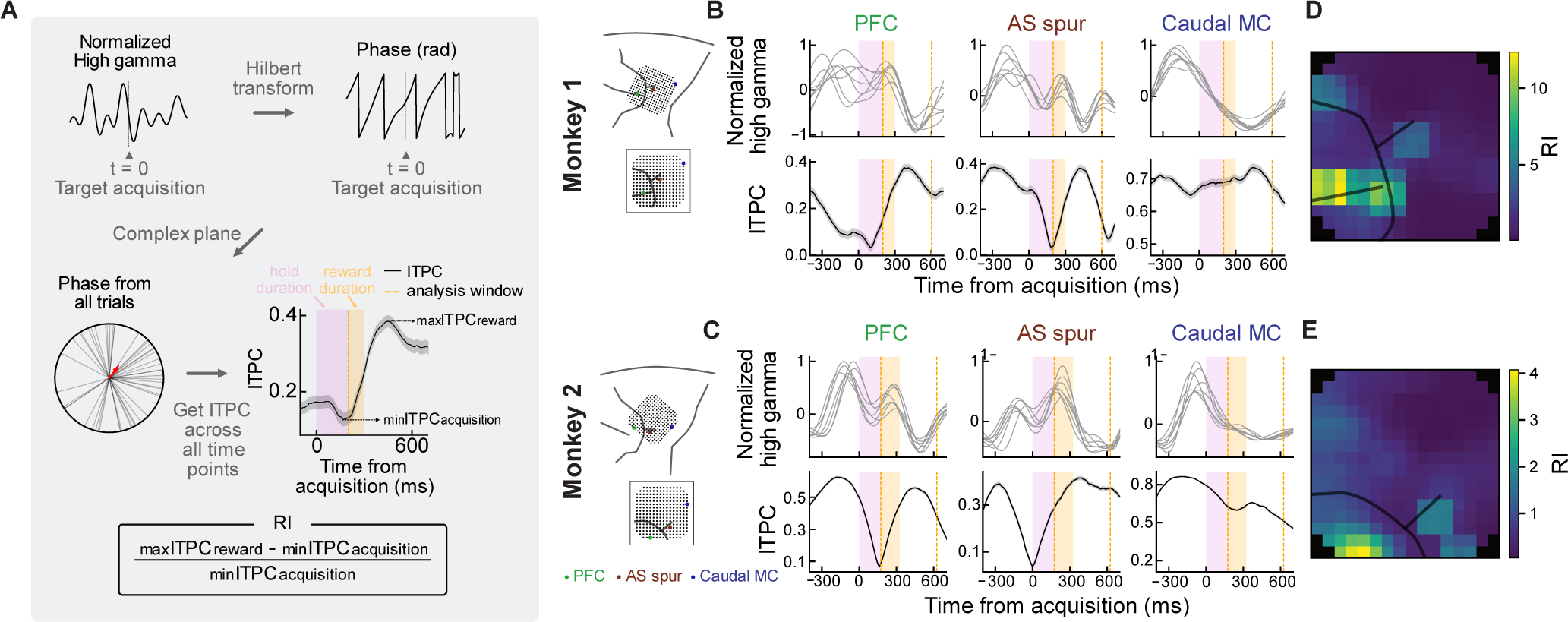
Reward information is localized within PFC and near AS in the motor cortex. **(A)** Schematic of the reward index (RI) analysis with data from an example channel for each calculation step. A Hilbert transform was applied to the filtered high-gamma power aligned to the target acquisition. Phases were averaged across all trials to obtain ITPC as a function of time, which was used to calculate the RI. **(B, C)** Filtered high-gamma power (top) and ITPC aligned to target acquisition (bottom) in example channels located in PFC, near AS, and in Caudal MC for monkeys 1 and 2, respectively. For high-gamma power, each line represents the mean power across trials separated by each target direction. For ITPC, solid lines indicate the mean and shaded areas indicate the standard deviations estimate from resampling. Magenta and orange shaded areas indicate the hold and reward durations, respectively. Orange dashed lines indicate the analysis window used to compute RI. Insets on the left show the array position with anatomical landmarks and indicate the location of example electrodes. **(D, E)** Spatial RI maps. Black lines indicate approximate sulci locations.

Examples of filtered high-gamma power, ITPC, and RI are depicted in Fig. 5B, 5C and Table 4. We observed two peaks of high-gamma activity before and after target acquisition in the channel over PFC and near AS. A dip in ITPC was sometimes observed during target acquisition, for example in the channels over PFC and near AS, but not in the channel over Caudal MC. RI was higher in the PFC channel and the near AS channel compared to the Caudal MC channel. Importantly, the heterogeneity in responses across channels suggests that these effects are unlikely to be related to physical recording artifacts that would be expected to impact all channels (e.g., licking, reward system signals).

**Table 4.** RI and CI in three representative channels.

| Monkey | Channel | RI | 95% CI |
| --- | --- | --- | --- |
| 1 | PFC | 22.7 | [5.35, 66.9] |
|  | AS spur | 23.2 | [5.77, 74.2] |
|  | Caudal MC | 0.0684 | [0.0296, 0.0997] |
| 2 | PFC | 7.11 | [5.54, 9.21] |
|  | AS spur | 9.15 | [6.19, 14.5] |
|  | Caudal MC | 0.248 | [0.225, 0.260] |

The spatial map of RI revealed highly localized reward information (Fig. 5D, E). Large RI was only observed within PFC and around the AS in PM for both monkeys. These results suggest that reward information, potentially reflecting reward detection and expectations, was clustered around the near AS in PM and within PFC. We also examined non-normalized RI maps (not shown), which also showed strongest reward indices within PFC and around the AS in PM, while also indicating presence of reward information (i.e. non-zero RI values) distributed across motor cortices.

## Discussion

Neural computations during goal-directed reaching behaviors are often studied in subsets of brain regions and in isolation from other task-relevant information such as eye movements and rewards. We used a reaching task with minimally or unconstrained eye movements to dissect eye movement, arm movement, and reward information across a significant portion motor and prefrontal cortices using µECoG. Functional maps of directional encoding varied depending on task events, illustrating the progression of movement information across the cortex. Functional maps not only contained spatially localized information about arm movement direction, but also eye movement direction and reward-related signals in high-gamma power. Our findings reveal more details of the functional organization of frontal motor areas and highlight challenges and opportunities for interpreting activity in these regions for applications like BCIs.

### Mapping changes in directional encoding across cortical areas

An advantage of surface electrode arrays is the ability to densely sample information across the cortical surface. µECoG revealed temporally varying spatial patterns of directional information (Fig. 3). Maps of both modulation depth and target decoding showed task-relevant activity moving from PFC, then rostral MC, and finally caudal MC. These temporal shifts across brain regions likely correspond to the sequence of behaviors in the task. The early presence of directional information in the PFC likely reflects visual processing or eye movement-related activity given that DLPFC and FEF receive low-latency visual information (30 – 150 ms) and contribute to saccade generation and control (Funahashi et al., 1990; Thompson et al., 1996). Rostral MC also exhibited earlier directional information, while Caudal MC exhibited directional information later during movement, consistent with a functional gradient from motor preparation to execution (Johnson et al., 1996).

We observed task-related modulation in multiple frequency bands, including delta and high-gamma (Fig. 2), consistent with past work (Rickert et al., 2005; Ball et al., 2009). Our decoding results are also consistent with observations that delta and high-gamma bands contain information about movement direction. This may be driven by high-gamma’s relationship to spiking activity (Ray et al., 2008) and delta’s relationship to low-frequency signals like the local motor potential (Schalk et al., 2007).

### Overlap of eye movement and arm movement information

Monkeys reached within tens of milliseconds of moving their eyes to the target (Fig 1). Analyzing the phase of high-gamma power (ITPC and EPI analysis) resolved differences in neural activity patterns across brain regions related to eye or arm movements. Our EPI maps suggested that PFC and MC primarily had eye and arm information, respectively, consistent with expectations from neuron recordings in each area (Georgopoulos et al., 1982; Funahashi et al., 1990).

Our ITPC analysis also revealed that the phase of high-gamma power around the AS spur in PM was aligned to saccade onset or both saccade and reach onsets, which implies that eye information and arm information were mixed. This finding is consistent with observations that some neurons in this region discharge for both eye and arm movement initiation (Kurata, 2017) with timing that is correlated with both saccade and reach onsets (Neromyliotis and Moschovakis, 2017). Our results highlight µECoG’s ability to resolve functional distinctions across brain regions and further suggest that activity in PM near the AS may contribute to generating coordinated eye and arm movements.

While ITPC analyses showed a bias towards arm information in MC, we also observed phase-relationships with eye movements in MC. Past work shows that MC neurons contain eye-related activity thought to reflect transformations between eye-centered and arm-centered coordinates (Pesaran et al., 2006; Batista et al., 2007). Consistent with this, our decoding analysis revealed a significant influence of free fixation on neural signals after reach onset in MC. Our results support the integration of eye and arm information in MC and map these interactions across the rostral-caudal expanse of MC.

### Reward information is localized near the principal sulcus and spur of the arcuate sulcus

Reward influences saccade onset, reach onset, movement variability, and learning (Takikawa et al., 2002; Izawa and Shadmehr, 2011; Manohar et al., 2015; Kojima and Soetedjo, 2017; Summerside et al., 2018). Reward-related activity has primarily been studied in individual frontal cortical regions. µECoG allowed us to map the spatial distribution of reward activity, which revealed focal reward-related regions in PFC and around the AS spur in PM (Fig. 5).

Within PFC, activity was primarily near the PS in both monkeys, though we had limited coverage of PFC in Monkey 2.

Our findings are consistent with past neuron recordings. For instance, DLPFC and PM near the AS exhibit reward-related activity during reward delivery even when the amount of reward is constant across trials (Watanabe, 1989; Ichihara-Takeda and Funahashi, 2006; Ramkumar et al., 2016). Our task kept the amount of reward consistent across trials. This suggests that reward-related activity in these regions may represent reward detection and expectations, as opposed to alternate computations like a reward prediction error.

The timing of phase-consistency in high-gamma power may provide clues as to the source of reward signals along the PS and AS. The latency of reward-related ITPC in high-gamma appears to be slower than phasic activity of dopamine neurons in the midbrain (130 ms, Ljungberg et al., 1992). This longer latency responses is consistent with neuron recordings in DLPFC and PMd, which estimate reward response times of 282.7 ± 125.1 ms and 400-600 ms, respectively (Ichihara-Takeda and Funahashi, 2006; Ramkumar et al., 2016). Dopamine neurons in the midbrain project to the striatum in the basal ganglia (Schultz, 2000), which is reciprocally connected with cortex (Middleton and Strick, 2000; Haber and Knutson, 2010).

Considering the slower latency in DLPFC and PMd, reward-related information may come from this multi-synaptic cortico-basal ganglia loop.

A recent study showed that neuronal activity in the motor cortex also contains information about reward prediction errors alongside arm movement information (Ramakrishnan et al., 2017). Although our task design did not allow us to investigate the details of reward representations, our results suggest that µECoG may help map details of reward encoding across cortical regions.

### Similarities and differences between monkeys

Many of our core findings were similar across monkeys, though our maps also revealed interesting differences. Spatiotemporal trends in maps were broadly consistent across monkeys, including: 1) patterns of directional information over time across brain regions (Fig. 3), 2) patterns in eye versus arm preference (PFC biased towards eye information, regions with both eye and arm information clustered around the AS spur, and biases towards arm information primarily in MC, (Fig. 4)), and 3) reward-related activity localized within PFC and along the AS spur in PM (Fig. 5). These similarities provide evidence that our µECoG measurements and analyses robustly resolved functional maps of neural processing.

While patterns across brain regions were consistent, we see clear differences across animals that may be related to chamber targeting and behavior differences. Monkey 2’s implant was shifted rostromedially compared to Monkey 1 which made us unable to record signals in the whole DLPFC in Monkey 2. This is likely why the EPI map from Monkey 2 displays less eye preference than Monkey 1 (Fig. 4). Importantly, our monkeys demonstrated different patterns of eye movements during the task. Even with eye position constraints, Monkey 2 often generated irrelevant saccades. Since saccades can influence neural signals (Fig. 4), these saccades likely influenced the functional maps (Fig. 3).

While we were able to better match the two monkeys’ behavior via altered task constraints in Monkey 2, this constraint may further contribute to differences between animals. Inhibiting eye movement in a saccade task has been shown to stop task-irrelevant arm movement (Wessel et al., 2013). Consistent with this observation, Monkey 2 showed slower reach onset in the CF condition than the FF condition (mean reach onset: 175 ms for FF, 259 ms for CF, *p* = 1.60 × 10^-51^, Wilcoxon rank sum test). This result may be related to an increased contribution of beta power to decoding in Monkey 2 (Fig 3O), since beta power is thought to relate to top-down signals reflecting movement inhibition in frontal cortices (Hwang et al., 2014; Khanna and Carmena, 2017; Barone and Rossiter, 2021). Requiring Monkey 2 to constrain his eye movements may have changed the computations required in the task (inhibiting movements), producing corresponding changes in neural signals (beta power) and functional maps. While we could have matched task constraints between monkeys, Monkey 1 primarily generated relevant saccades after the go-cue in the free fixation task (0.052% of his trials had a saccade before the go-cue). Thus, a controlled fixation task would be unlikely to impose a significant demand to inhibit eye movements for Monkey 1. Our findings highlight natural variability in how animals perform behavioral tasks, nuances in using task demands to shape behavior, and the importance of monitoring eye movements in visually guided reaching tasks.

### Implications for brain-computer interfaces

µECoG offers an alternative to penetrating electrodes that has potential for long term signal stability (Chao et al., 2010), which may have advantages for BCIs (Flint et al., 2013; Benabid et al., 2019; Silversmith et al., 2021). Our study demonstrates that both eye and arm movement information can be detected from µECoG signals within motor cortices (Fig. 3 and 4), which presents challenges and opportunities. Decoding movement information from two effectors may enhance the versatility of BCIs, but it also requires decoding algorithms that can distinguish between these distinct motor signals.

Current BCIs often rely heavily on visual feedback and operate in controlled settings, which likely leads to closely correlated eye and arm movements. We found that ability to decode movement targets from both rostral and caudal MC improves when eye movements are restricted (Fig. 4), showing that unconstrained eye movements can confound arm movement decoding. This aligns with prior findings that incorporating knowledge of gaze location can improve arm movement decoding from neurons in PMd (Batista et al., 2008), and extends this finding to µECoG signals across much of MC. These findings underscore the need for BCI algorithms that are robust to eye movement variability, which may require developing approaches to isolate eye and arm information. The presence of eye signals in motor areas also opens the opportunity to leverage this information to enhance decoding. For instance, recent work suggests that eye movements provide useful training signals to dynamically update decoding algorithms for robust and generalizable performance (Chou et al., 2024).

The presence of reward-related activity in motor cortices (Fig. 5) similarly presents challenges and opportunities for BCI. Approaches to isolate reward information may be needed for robust movement decoding. Yet, neural signals related to task errors have also been used to guide BCI decoder training (Mahmoudi and Sanchez, 2011; Rouanne et al., 2022). BCIs could leverage reward information to adapt algorithms, adjust task difficulty, and so on.

Our results shed further light on the richness of neural activity in frontal motor cortices, revealing distributed processing across brain regions reflecting multiple aspects of behavior during reaching. Future work to disentangle these computations will improve our understanding of motor control and improve our ability to build robust, personalized BCIs.

## Acknowledgments

We would like to thank Jonathan Viventi and colleagues for providing the electrode arrays used in this work, and the Washington National Primate Research Behavioral Management team for their consultations on animal training. This work was supported in part by a Nakajima Foundation fellowship (T.O.), a postdoctoral fellow award from Weill Neurohub (L.R.S.), an NSF Accelnet INBIC fellowship (P.R.), a National Center for Advancing Translational Sciences of the National Institutes of Health fellowship (TL1 TR002318, R.A.C.), a Simons Collaboration for the Global Brain Pilot award (898220, A.L.O.), the Eunice Kennedy Shriver National Institute of Child Health and Human Development (NIH K12HD073945, A.L.O.), the National Institute of Neurological Disorders and Stroke (NIH R01 NS134634, A.L.O.), and the NIH Office of Research Infrastructure Programs (P51 OD010425).

